# Pipeline for species-resolved full-length16S rRNA amplicon nanopore sequencing analysis of low-complexity bacterial microbiota

**DOI:** 10.1101/2023.12.05.570138

**Authors:** Disha Tandon, Yuan Dong, Siegfried Hapfelmeier

**Author notes:** Address correspondence to corresponding author Siegfried Hapfelmeier and co-corresponding author Disha Tandon.

## Abstract

16S rRNA amplicon sequencing is a fundamental tool for characterizing prokaryotic microbial communities. While short-read 16S rRNA sequencing is a proven standard for examining complex microbiomes, it cannot achieve taxonomic classification beyond genus level. Third-generation sequencing technologies, particularly nanopore sequencing, have allowed for full-length 16S rRNA gene sequencing enabling enhanced taxonomic resolution to species and strain levels.

Precise species-to-strain level classification is crucial in investigating low-complexity microbiota. This research presents an efficient pipeline using full-length 16S rRNA amplicon nanopore sequencing, spanning library prep to computational analysis for low-complexity microbiota composition analysis. We applied this pipeline to a defined intestinal bacterial community in gnotobiotic mice to evaluate different methods.

Our findings revealed that the proprietary barcoded universal primers 27F-1492R from Oxford Nanopore Technologies(ONT) 16S amplicon sequencing kit did not efficiently amplify the 16S rRNA gene of *Bifidobacterium* species. Addressing this constraint, we designed degenerate primers and employed ONT’s native barcoding kit for library preparation. We developed a customized wet lab and bioinformatics pipeline for processing and classifying amplicon reads at the species level.

Validation of the protocol using a mock community DNA sample with known composition confirmed a reduced analytical bias. Additionally, our method surpassed Illumina short-read V3-V4 amplicon sequencing, achieving accurate species-level classification compared to Illumina’s genus-level accuracy.

This pipeline is tailored for analyzing the composition of low-complexity microbiota from natural ecosystems and synthetic/gnotobiotic communities. It is cost- and time-effective and therefore accessible for small-scale studies that would otherwise be hindered by the typically long turnaround times of NGS services.

**Importance:** 16S rRNA amplicon sequencing is conventionally used to identify microbes and determine their composition in microbial communities. Deep amplicon sequencing of complex microbiomes is well established using short-read sequencing targeting variable regions of the 16S rRNA gene. Short reads enable the classification of bacteria until the genus level in the taxa hierarchy, whereas long reads provide better chances of identifying bacteria to species and even strain levels. This study introduces a streamlined approach for analyzing simple microbial communities using full-length 16S rRNA amplicon nanopore long read sequencing. This approach surpasses Illumina sequencing in species accuracy, is cost-effective and time-efficient. Tailored for low-complexity microbiota, it facilitates studies in natural or synthetic communities, especially beneficial for smaller-scale projects with limited resources.

## Introduction

Low-complexity microbiota, characterized by low microbial species diversity, is found in various natural ecosystems, such as the adult bee gut, marine sponges, and pesticide-affected and nutrient-depleted soils^1–4^. In parts of the human gastrointestinal microbiota, low complexity is observed in specific locations (e.g. in the stomach mucosa) or temporarily (e.g. in the small intestine in early life), while still displaying high interindividual diversity^5–7^. Additionally, low-complexity microbiomes in the form of “synthetic microbial communities” (SynComs) of defined strain composition are valuable experimental tools for studying host-microbe and microbe-microbe interactions in gnotobiotic animals and plants, or *in vitro* model systems^8–11^. Besides their apparent simplicity, the unbiased compositional analysis of such microbial communities presents a specific challenge for current untargeted DNA sequencing based methods.

Among current sequencing-based methods for microbial compositional analysis, 16S rRNA gene amplicon sequencing is the most well-established and widely used approach for inferring relative abundance patterns^12–14^. The 16S rRNA gene, conserved in all prokaryotes, evolves slowly compared to most other regions of the prokaryotic genome. It contains hypervariable regions (V1-V9), which recapitulate most of the phylogenetic diversity among bacteria and archaea. These regions can be selectively PCR amplified using universal primers (that are complementary to lowly variable regions flanking one or several V regions), and the resulting sequences analyzed using high-throughput sequencing technologies. Computational tools are used to taxonomically classify and calculate the relative abundances of 16S rRNA gene sequence reads, inferring microbial composition. Short-read technologies like Illumina sequencing, despite their limitations in read lengths (up to 2*251 bp for paired-end sequencing, a fraction of the full 16S rRNA gene’s ∼1500 bp), offer high fidelity and can resolve single-nucleotide sequence variants within individual or paired consecutive hypervariable regions. This capability enables classification down to phylum, class, order, family, and sometimes even genus level. However, for reliable genus- and species-level classification, the sequencing of full-length 16S rRNA genes or multiple (>2) hypervariable regions is typically necessary^15–18^. In previous studies, *in silico* methods have emphasized the advantages of longer sequences over shorter ones for bacterial compositional analysis^16,17^. Additionally, it has been shown that different combinations of V-regions result in different compositional data, potentially introducing analytical bias.

Illumina has been the leading next-generation sequencing (NGS) technology for short-read sequencing of 16S rRNA amplicons spanning one or more hypervariable regions. Among these, Illumina’s short-read sequencing technology remains state-of-the-art due to its low sequencing error rate, achieving a base call accuracy of 99%. Nevertheless, the majority of NGS technologies entail cost- and labour-intensive processes, in addition to requiring high-end sequencing equipment and laboratory resources. As a result, most academic laboratories now rely on either large-scale sequencing core facilities within academic institutions or outsource their sequencing needs to commercial providers, which offer competitive services. However, one significant limitation remains for large-scale NGS, and that is the sample-to-data turnaround times that typically span weeks to months. This hinders applications in diagnostics and experimental designs that necessitate rapid microbiome analyses. Furthermore, laboratories in low-resource settings often face limited access to large-scale NGS sequencing.

In recent years, long-read sequencing technologies have gained popularity for their ability to directly sequence DNA molecules spanning several kilobases. These technologies, pioneered by Oxford Nanopore Technologies (ONT) and Pacific Biosciences (PacBio), are commonly referred to as third-generation sequencing technology. So far, both ONT (6%-8% error rate) and PacBio (up to 13% error rate, with up to 99.999% base call accuracy achievable through multiple-pass sequencing) exhibit relatively high error rates^19,20^. However, the advantage of long-read sequencing lies in its extended read length, leading to increased overall alignment quality compared to short reads^21^. Long-read sequencing can provide high alignment quality even at low sequencing depths, while short-read sequencing typically requires high sequencing depth for optimal alignment. PacBio long-read sequencing remains cost-prohibitive for widespread adoption in settings with limited resources. In contrast, ONT sequencing instruments and chemistry are highly affordable, user-friendly, and efficient, making complex sequencing experiments accessible even on a small scale^22^. ONT long-read sequencing offers high coverage of the full-length 16S rRNA gene, reducing the likelihood of misaligned highly conserved regions, despite its relatively high error rate. Many researchers have consequently started to employ long-read ONT 16S rRNA amplicon sequencing for microbial compositional analysis^12,23–26^.

ONT offers a combined PCR amplification-barcoding kit (16S Barcoding Kit 1-24, product number SQK-16S024) designed for efficient multiplexed full-length 16S rRNA amplicon sequencing. A streamlined protocol combines the amplification and barcoding of the full-length 16S rRNA gene in a single PCR step. ONT proprietary amplification primers feature the universal 27F and 1492R priming sequences along with barcode sequence tags. The universal 16S rRNA gene primer sequences 27F and 1492R have been well-established as specific and sensitive for PCR amplification of diverse bacterial 16S rRNA gene sequences in various sequencing-based microbial identification and quantification methods^27–32^. However, some prior studies have noted the low specificity of these primers for *Lactobacillales* and *Bifidobacterium* species isolated from infant human gut^5,33^. Thus, when studying microbial communities that include these taxa, a revised primer design may be more suitable. As of the time of writing, ONT provides native barcoding protocols with kits like NBD-104, NBD-114, and NBD196, allowing the attachment of ONT sequencing barcodes to PCR amplicons generated with user-defined primers and PCR protocols.

In this study, we confirm previous reports of suboptimal performance of primer pair 27F/1492R ^5,33^. We demonstrate that library preparation and sequencing with the ONT SQK-16S024 kit consequently results in inaccurate classification of gnotobiotic mouse intestinal bacterial strains. To address this, we have designed optimized degenerate primers and utilized amplicon multiplexing by native barcoding to show their applicability and reduced bias.

While several bioinformatic tools exist for analyzing nanopore sequencing data, an open-source approach with high sensitivity and specificity has been lacking^34–37^. To address this gap, we developed a bioinformatics pipeline for processing long-read sequences generated by nanopore 16S rRNA gene amplicon sequencing and classifying them to appropriate taxa at approximately 88% similarity level. This integrated method, from library preparation to bioinformatic analysis, is optimized for the analysis of low-complexity bacterial communities, such as gnotobiotic mouse gut microbiota, by nanopore amplicon sequencing. Benchmarking against Illumina V3-V4 amplicon sequencing confirms that, as demonstrated by others^16,38^, long read amplicon sequencing, compared to short-read-based approaches, reduces misalignment of reads between phylogenetically similar taxa.

## Methods

### Animal experiments and gnotobiology

All animal experiments were approved by the Bernese Cantonal Ethical committee for animal experiments and carried out according to Swiss Federation law for animal experimentation under license BE66/19.

All animal experiments were performed with OligoMM12 associated gnotobiotic C57BL/6J mice^39^ that had been originally generated at the Clean Mouse Facility (CMF, University of Bern) by inoculating germ-free C57BL/6J mice with pure cultures of the OligoMM12 bacteria^39,40^ and had been stably maintained in flexible film isolators under strict bioexclusion. All animals studied are descendants of the same OligoMM12 mouse line and were housed in the Clean Mouse Facility (CMF) at the University of Bern. Mice were maintained at an ambient temperature of 23-25 °C and relative humidity of 52-60 %. All mice received a standard rodent chow diet (Kliba 3307, Ssniff, Germany) and autoclaved surgery irrigation water (Baxter, USA) *ad libitum*.

The OligoMM12 community comprises twelve murine intestinal bacterial species representative of five gut bacterial phyla that are naturally prevalent in the murine gastrointestinal tract^39^. The OligoMM12 community colonizes the mouse intestinal tract stably over multiple host generations^39–41^. The gnotobiotic mice studied to generate the data shown in Figure were OligoMM12 mice additionally colonized for 4 days with pure cultures of *Clostridium scindens* ATCC 35704 (a gift from Nigel Minton, University of Nottingham) and *Escherichia coli* MT1B1 (DSM 28618, DSMZ, Germany).

### Mock community metagenome preparation

9 of the OligoMM12 strains (listed in **Table 2**) were re-isolated from gnotobiotic OligoMM12 mice (Clean Mouse Facility, University of Bern) and cryopreserved. Each strain was grown individually on Columbia sheep blood (CSB; Oxoid, UK) agar under anaerobic conditions (Whitley A45 HEPA anaerobic station, Don Whitley Scientific, UK) at 37°C for a duration of 3 to 7 days. Single colonies of each individual strain were verified by MALDI-TOF analysis (Bruker Microflex LT) and subcultured anaerobically in liquid *Akkermansia* broth^41^ at 37 °C. Bacterial cells were harvested by centrifugation at 4800 rpm for 7 minutes at room temperature, and genomic DNA was extracted using the DNeasy PowerSoil Pro kit (Qiagen, USA). At this step, the strain identities were re-confirmed by 16S rRNA gene PCR amplification (using primer pair: 27F deg-1492R deg) followed by Sanger sequencing (Microsynth, CH). Mock community DNA was prepared by mixing equal concentrations of genomic DNA (determined using Qubit dsDNA BR or HS kits; Life Technologies, USA) of each bacterial strain. Three replicates mixed individually were used for subsequent analysis.

**Table 1.**
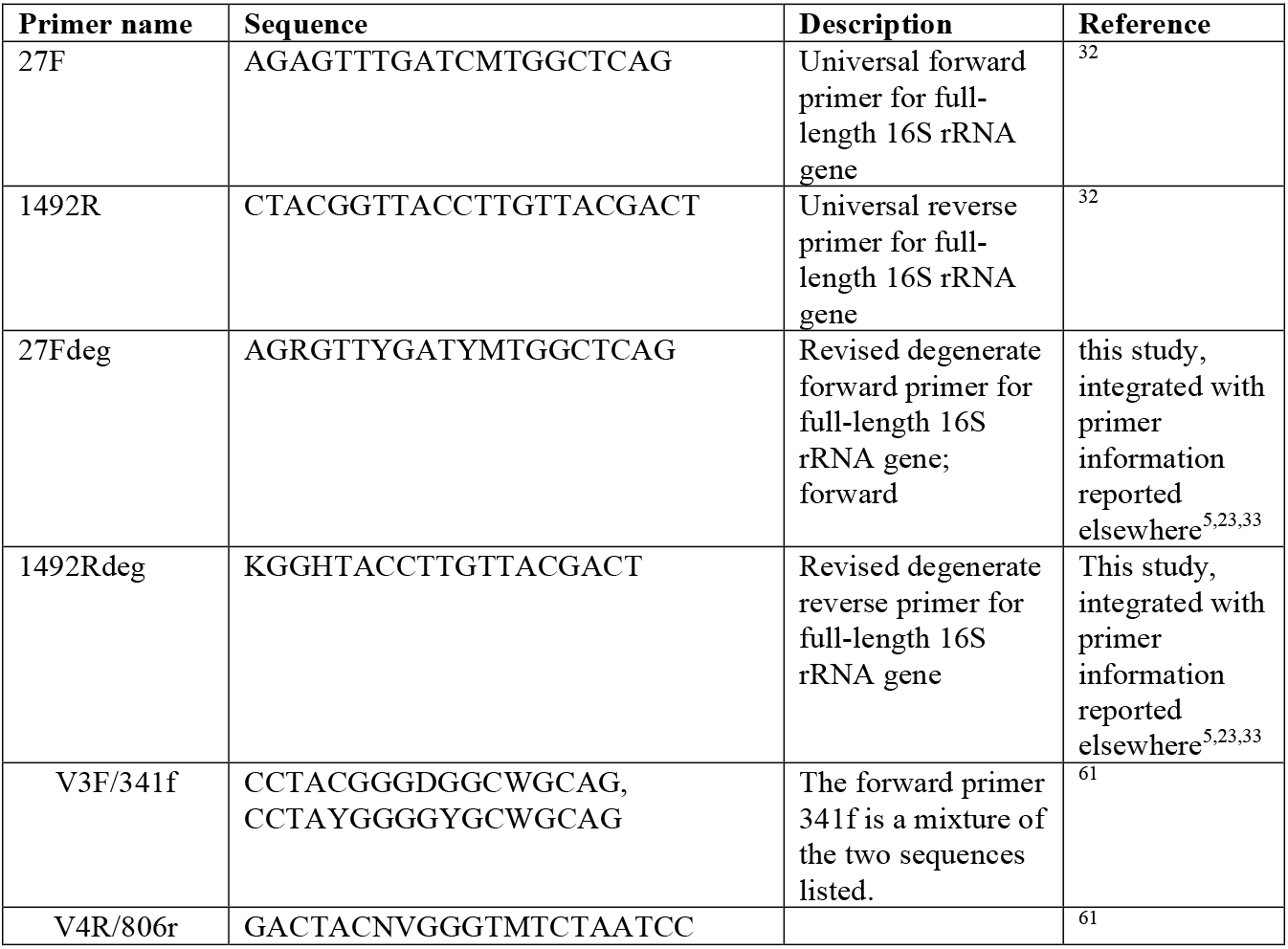
Primer sequences.

**Table 2.** Bacterial strains used in the mock community.

| Bacterial strain | Reference |
| --- | --- |
| *Bacteroides caecimuris* I48 | <sup>56</sup> |
| *Blautia pseudococcoides* YL58 | <sup>56</sup> |
| *Turicimonas muris* YL45 | <sup>56</sup> |
| *Enterococcus faecalis* KB1 | <sup>56</sup> |
| *Clostridium innocuum* I46 | <sup>56</sup> |
| *Flavonifractor plautii* YL31 | <sup>56</sup> |
| *Enterocloster clostridioformis* YL32 | <sup>56</sup> |
| *Limosilactobacillus reuteri* I49 | <sup>56</sup> |
| *Bifidobacterium animalis* YL2 | <sup>56</sup> |

### DNA extraction

Fresh intestinal content samples (feces content) from mice were collected in a 1.5 ml Eppendorf tube and resuspended in 1 ml sterile PBS. Metagenomic DNA was extracted using two kits (a) QIAmp Fast DNA Stool Mini Kit (Qiagen, USA) and (b) DNeasy PowerSoil Pro kit (Qiagen, USA). With the QIAmp Fast DNA stool kit, the protocol was modified to include an additional step of bead beating for 10 minutes at 50 Hz (TissueLyser LT, Qiagen, USA) with the PowerBead Pro tubes containing glass beads of diameter 0.5 mm (Qiagen, USA) before adding the lysis buffer as mentioned in the kit protocol. This step was added to enhance the homogenization of the sample.

### Quality control and Concentration

The quality control of the genomic DNA and the PCR DNA was done using Nanodrop (Thermo Fisher Scientific, USA) 260/230 and 260/280 ratios. The concentration of all DNA samples was determined by Qubit dsDNA BR or HS kits (Life Technologies, USA) on the Qubit 4.0 fluorometer. The quantity, purity and length of the total genomic DNA were assessed additionally on the Agilent FEMTO Pulse System with a Genomic DNA 165 kb Kit (Agilent, USA).

### Full-length 16S rRNA amplicon ONT sequencing using 16S-SQK024 kit

16S barcoding PCR barcoding was performed using the Oxford Nanopore Technology (Oxford Nanopore Technologies, UK) kit SQK-16S024. The primers provided in the kit include the unique barcode sequence attached to the universal 16S forward (27F) and reverse (1492R) primers (**Table 1**). The primers harbor the ONT proprietary rapid attachment chemistry required for adapter ligation. For the PCR, LongAmp Hot Start PCR Master Mix (New England Biolabs, USA) was used, which took approximately 1.5 hours to complete. The PCR was followed by gel electrophoresis to check the optimal amplification of ∼1500 bp fragments. Once the amplification was confirmed, the PCR product was purified using HighPrep PCR beads (Magbio Genomics, USA) on the DynaMAG.96 Side skirted Magnetic plate (Thermo Fisher Scientific, USA). Final Library preparation for the sequencing was performed using the recommended protocol for ONT kit SQK-16S024 (Oxford Nanopore Technologies, UK).

### Full-length 16S rRNA amplicon ONT sequencing using native barcoding kits

PCR amplification of the 16S rRNA gene was performed using either universal 27F-1492R primer pair or modified degenerate universal primers 27Fdeg-1492Rdeg (**Table 1**). PCR conditions include initial denaturation at 95° C for 5 minutes, followed by 30 thermal cycles of denaturation at 94° C for 30 seconds, annealing at 58° C for 30 seconds and elongation at 72° C for 40 seconds. This was followed by final end product extension at 72° C for 10 minutes.

PCR was followed by gel electrophoresis to confirm the amplification. PCR product was purified using MiniPrep PCR purification kit (Qiagen, USA). PCR purified product was end-prepared according to protocols of native barcoding kits (SQK-NBD-104, SQK-NBD-114 from Oxford Nanopore Technologies, UK) according to manufacturer’s instruction with minor modifications. The amplicons were purified using HighPrep PCR beads (Magbio Genomics, USA) on the DynaMAG.96 Side skirted Magnetic plate (Thermo Fisher Scientific, USA). The purified amplicons were barcoded using the native ONT barcoding kits SQK-NBD104/SQK-NBD114, according to manufacturer’s instructions, with the exception of increased starting amounts of amplicon DNA (200 ng/amplicon). The barcoded amplicon DNAs were pooled in equal concentrations and purified using HighPrep PCR beads on the DynaMAG.96 Side skirted Magnetic plate (Thermo Fisher Scientific, USA). In the final step, adapter ligation was carried out according to SQK-NBD104/SQK-NBD114 and SQK-LSK109 kit. Ligation was followed by processing the library for loading according to recommended protocol for SQK-LSK109 kit (Oxford Nanopore Technologies, UK).

Sequencing runs were performed on the R9.4.1 flow cell (Oxford Nanopore Technologies, UK).

### Illumina short read amplicon sequencing

NGS libraries targeting the V3-V4 region of the 16S rRNA gene were constructed using the Quick-16S Plus NGS Library Prep Kit (V3-V4; primer sequences in **Table 1**) (Zymo Research, USA) according to the instruction manual version 1.2.1. Two ng of input DNA was used for each sample, a BIO-RAD CFX96 Real-time System and C1000 touch thermal cycler were used for the qPCR-based steps. Alongside the samples, two negative controls were included; one with the buffer used to dilute the samples and one with the PCR mastermix lacking any DNA. Furthermore, a Microbial Community DNA Standard and a Microbial Community DNA standard II (log distribution) were also processed (Zymo Research, USA, respectively). The final library pool was evaluated using a Thermo Fisher Scientific Qubit 4.0 fluorometer with the Qubit dsDNA HS Assay Kit (Thermo Fisher Scientific, USA) and an Agilent Fragment Analyzer (Agilent, USA) with HS NGS Fragment Kit, respectively. The library pool, including all controls, was paired-end sequenced using a MiSeq Reagent Kit v3 600 cycles (Illumina, USA) on an Illumina MiSeq instrument. The quality of the sequencing run was assessed using Illumina Sequencing Analysis Viewer (Illumina version 2.4.7) and all base call files were demultiplexed and converted into FASTQ files using Illumina bcl2fastq conversion software v2.20. All steps post gDNA extraction to sequencing data generation and data utility were performed at the Next Generation Sequencing Platform, University of Bern, Switzerland.

### Bioinformatic analysis

#### Nanopore

Samples with more than 1000 reads were considered for further analysis. Base-called raw reads were obtained from all runs using the ‘guppy’ basecaller^42^. Demultiplexing was performed using ‘qcat’ with parameters enabled for trimming the adapters^43^. Chimeras were checked by mapping the reads against all reads using ‘minimap2’^44^ and then detecting the chimeric reads using ‘yacrd’^45^. The reads were filtered by quality (>=9) and length (1000-1600) using ‘nanofilt’^46^. For each sample, reads were normalized by the copy number of the 16S rRNA gene in the respective bacterial genome and by the total number of reads sequenced. Reads were aligned and classified using ‘EMU’^47^ against a custom database that includes sequences from rrnDB v5.6^48^ and NCBI 16S RefSeq^49^ databases. EMU aligns the reads against the reference database using minimap2 and uses an expectation-maximization approach to generate taxonomic abundance values. Results were plotted through in-house R scripts using ggplot2 package^50^.

#### Illumina

Samples with more than 80000 reads were considered for further analysis. Low quality bases (10 bp at the ends) and adapters were trimmed from the demultiplexed reads using cutadapt^51^. Ribosomal Database Project (RDP) Classifier was used for classifying the reads against the RDP database^52^. Reads with classification until the genus taxon was considered for interpretation. Results were plotted through in-house R scripts using ggplot2 package^50^.

#### Other analysis

Multiple sequence alignment was performed using the ‘mafft’ tool^53^ and visualized in Jalview^54^. Euclidean distance was calculated using the following formula through an in-house R script.

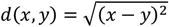

Where *d(x,y)* refers to distance between two vectors/points *x* and *y*.

## Data availability

Nanopore and Illumina sequencing data generated in this study is available at the European Nucleotide Archive (Accession num: PRJEB70478).

## Results

### Evaluation of ONT proprietary native barcoding protocol

The OligoMM12 community comprises of twelve murine intestinal bacterial species representative of five gut bacterial phyla that are naturally prevalent in the murine gastrointestinal tract and had been isolated from laboratory mice and fully sequenced^39^. The OligoMM12 community colonizes the intestinal tract of gnotobiotic mice stably over multiple host generations^39–41^. Previous work has characterized short-term and long-term compositional dynamics of OligoMM12 in gnotobiotic mice in detail using short-read amplicon and shotgun sequencing or 16S rRNA gene specific qPCR^40^. Several studies have evaluated the reproducibility of OligoMM12 composition between animal facilities, its vertical transmission and early life development, and response to diet shifts^39–41,55^.

We first evaluated ONT sequencing of full-length 16S rRNA gene amplicons generated from DNA extracted from fecal contents of adult 6-8-week-old OligoMM12 mice using universal primer pair 27F/1492R (**Table 1**). Barcoding, subsequent library preparation, and sequencing were carried out according to manufacturer’s instructions (ONT native barcoding kits NBD104/114 and LSK109). Using this protocol, we were able to corroborate the previously published information that the most highly relatively abundant strain in the fecal microbiome of gnotobiotic OligoMM12 mice maintained on standard rodent chow diet is *Bacteroides caecimuris* I48^39,40^ (**Figure 1a**). Moreover, as reported by others, *Bifidobacterium animalis* YL2, known to be lowly abundant in the feces of adult mice, could not be detected^40,41^ (**Figure 1a**).

**Figure 1.**
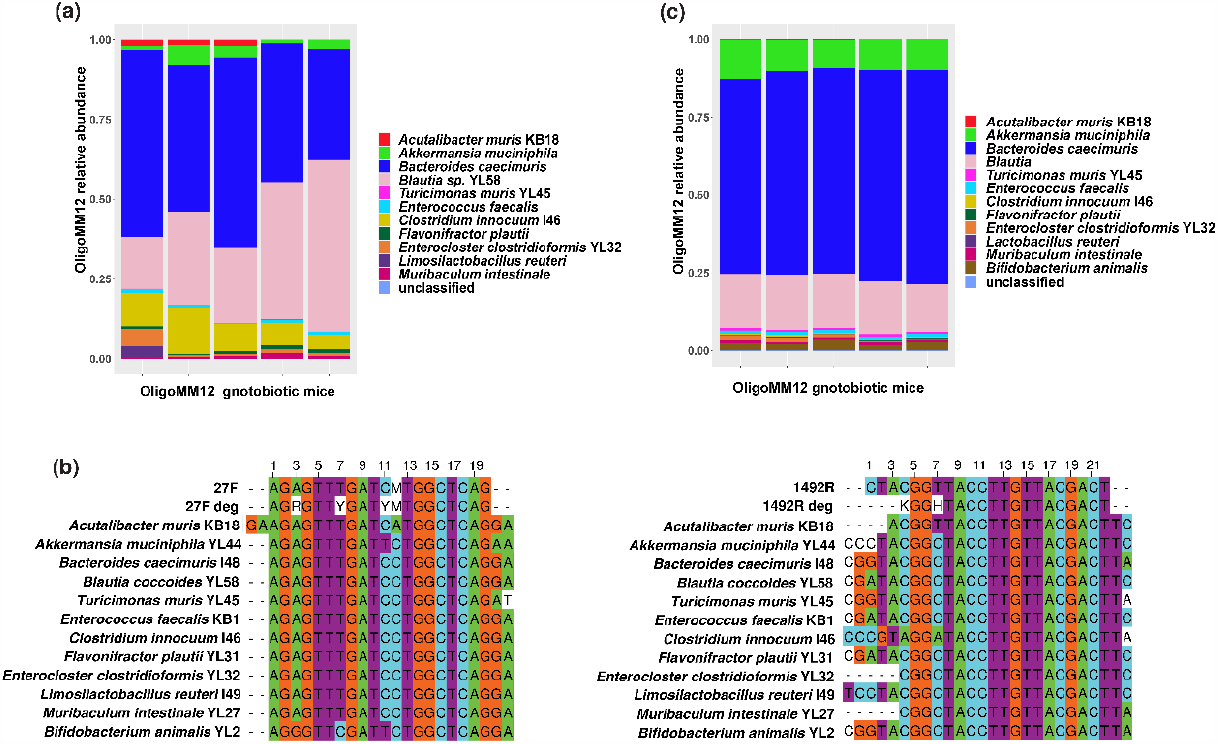
Degenerate primer pair 27Fdeg-1492Rdeg amplifies 16S rRNA genes from all bacterial species of OligoMM12 mice. Bacterial compositional analysis of fecal contents from OligoMM12 mice was performed by full-length 16S rRNA gene amplicon sequencing using **(a)** universal primer pair 27F-1492R, and **(c)** revised degenerate universal primer pair 27Fdeg-1492Rdeg, respectively. Sequencing libraries were prepared using ONT native barcoding kit (SQK-NBD104/SQK-NBD114). Legend indicates the taxonomic levels (species and strain) to which the reads could be classified. Reads with strain level classification are indicated with strain numbers (KB18, YL58, YL45, I46, YL32) in the legend. **(b)** Multiple sequence alignment of primer pairs 27F-1492R and 27Fdeg-1492Rdeg with the flanking ends of the 16S rRNA genes of OligoMM12 community members. Degenerate bases - R = A/G, Y = C/T, M = A/C, K = G/T, H = A/C/T.

### Revised degenerate primer design for detection of lowly abundant taxa

In a prior study employing 16S rRNA gene-specific qPCR, a marked reduction in the absolute abundance of *Bifidobacterium animalis* YL2 was noted during the initial ten days of *de novo* colonization of adult germ-free mice with pure culture of OligoMM12 bacteria^39^. Additionally, Yilmaz and colleagues, utilizing short-read shotgun metagenomic sequencing, reported that strain YL2 was only detectable only in young (2-3 weeks old), but not in adult OligoMM12 mice^40^. However, others have shown that the universal primer pair 27F/1492R lacks sequence complementarity for Lactobacillales and *Bifidobacterium* species^5,33^. This led to our hypothesis that protocols based on 27F/1492R universal primers may underestimate or entirely fail to quantify the relative abundance of strain YL2. We performed a multiple sequence alignment of flanking regions of 16S rRNA full length sequences of all 12 OligoMM12 bacterial strains obtained from NCBI genbank (**Figure 1b**). We found 3 mismatches between forward primer 27F and the YL2 genomic 16S priming site at positions 3,7, and 11 (**Figure 1b**). We additionally found 2 mismatches between reverse primer 1492R and the reverse priming site in YL2 (positions 1 and 7), as well as 4 mismatches in I46 (positions 2,3,4, and 7) and 1 mismatch (position 7) in all remaining OlioMM12 strains except KB18 (**Figure 1b**). Based on this consensus alignment information we designed the revised degenerate primer pair 27Fdeg/1492Rdeg (see **Table 1**) that is optimized according to predicted sequence complementarity. We then tested the revised primer pair in combination with the ONT native barcoding kits (NBD104/114 and LSK109) for full-length 16S rRNA gene amplicon sequencing analysis. Supporting our hypothesis, the modified protocol consistently enabled the quantification of low-abundance *Bifidobacterium animalis* YL2 in the cecum of adult OligoMM12 animals (**Figure1c**). This underscores the suitability of the revised universal primer pair 27Fdeg/1492Rdeg for *Bifidobacterium* quantification.

### PCR amplification followed by native barcoding yields more accurate taxonomic profiling compared to the 16S-SQK024 protocol

ONT’s proprietary direct multiplexing 16S rRNA gene amplicon kit 16S-SQK024 offers a streamlined workflow based on the standard 27F/1492R universal primers. To assess potential biases expected from inefficient primer design, we conducted a systematic comparison of native ONT barcoding using the optimized 27Fdeg/1492Rdeg primers.

In this comparison, we included a mock community DNA control, created by mixing genomic DNAs from 9 individually cultured OligoMM12 strains (**Table 2**) in equal concentrations. This mock-community (in three replicates) and metagenomic DNA extracted from the fecal microbiota of gnotobiotic mice colonized with 13 (the OligoMM12 community and *Escherichia coli* MT1B1^56^) or 14 (OligoMM12, *Escherichia coli* MT1B1, and *Clostridium scindens* ATCC35704^56^) bacterial species were analyzed using two approaches: Approach 1, utilizing ONT kit 16S-SQK024 combining amplification and barcoding through 27F/1492R primers (**Table 1, Figure 2a**), and Approach-2, utilizing our re-designed degenerate primers 27Fdeg/1492Rdeg for amplification (**Table 1**), followed by ONT native barcoding through ONT kits NBD104/NBD114 (**Figure 2b**).

**Figure 2.**
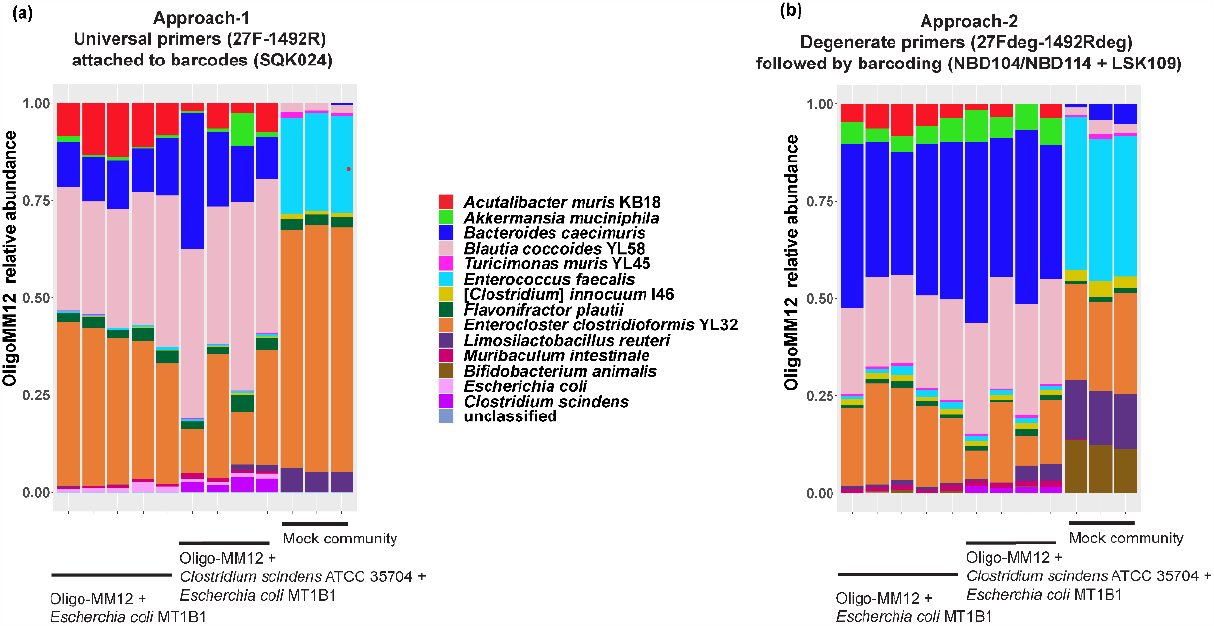
Comparison of two ONT library preparation approaches. **(a)** Approach-1, using ONT kit SQK-16S024, including proprietary pre-barcoded amplification primers based on universal primers 27F-1492R. **(b)** Approach-2, 16S gene amplification using revised primers 27Fdeg/1492Rdeg followed by multiplex barcoding using ONT kits SQK-NBD104/SQK-NB114 and SQK-LSK109 kit). Legend indicates the taxonomic levels (species and strain) to which the reads could be classified. Strain-level classifications are indicated by strain designations (KB18, YL58, YL45, I46, YL32); species-level classifications are shown as species name only. Each stacked bar represents one mouse individual or mock-community replicate, displayed in the same order in both panels.

To evaluate sequencing analysis bias, we calculated the Euclidean distance between the expected mock-community genomic composition, and the microbial compositional results obtained using either sequencing protocol. The Euclidean distance metric provides a straightforward way to calculate the similarity between predicted and observed microbial composition at the bacterial species level (see Methods section for details). The higher the distance, the more is the variation, and the lower the distance, the less is the variation. The mock community comprised 9 bacterial strains at equal abundance (0.111 expected relative abundance each). The observed relative proportions were derived from the relative abundance data obtained from the sequencing data analysis. **Table 3** summarizes the Euclidean distances between expected and observed relative abundances of each mock community member for both sequencing approaches A1 and A2. On average, Approach-1 exhibited higher distances (0.587) than Approach-2 (0.357), suggesting that Approach-2 is overall less biased (see **Table 3** and **Figure 2**). Among individual bacteria, the highest Euclidean distance (0.513) in Approach-1 was observed for *Enterocloster clostridioformis* YL32, indicating its overrepresentation (**Figure 2**). The highest individual distance in Approach-2 was calculated for *Enterococcus faecalis* (0.261; **Figure 2**), suggesting overrepresentation. The distance for *Bifidobacterium animalis* was approximately 10-fold higher in Approach-1 (0.111) compared to Approach-2 (0.014), indicating reduced efficiency in detection for Approach-1. Overall, our findings demonstrate that Approach-2, utilizing modified degenerate primers 27Fdeg/1492Rdeg and native barcoding with the NBD104/NBD114 kit, better represented the gnotobiotic community (**Figure 2**).

**Table 3.** Euclidean distance of expected and observed mock community microbial composition.

| Bacterial strain | Approach-1 |  |  |  | Approach-2 |  |  |  |
| --- | --- | --- | --- | --- | --- | --- | --- | --- |
|  | R1 | R2 | R3 | Average | R1 | R2 | R3 | Average |
| *Bacteroides caecimuris* | 0.111 | 0.111 | 0.107 | 0.110 | 0.102 | 0.071 | 0.061 | 0.078 |
| *Blautia pseudococcoides* YL58 | 0.088 | 0.092 | 0.090 | 0.090 | 0.093 | 0.072 | 0.087 | 0.084 |
| *Turicimonas muris* YL45 | 0.096 | 0.105 | 0.103 | 0.102 | 0.104 | 0.099 | 0.103 | 0.102 |
| *Enterococcus faecalis* | 0.135 | 0.142 | 0.137 | 0.138 | 0.281 | 0.252 | 0.249 | 0.261 |
| *Clostridium innocuum* I46 | 0.099 | 0.103 | 0.099 | 0.100 | 0.083 | 0.069 | 0.079 | 0.077 |
| *Flavonifractor plautii* | 0.079 | 0.083 | 0.084 | 0.082 | 0.104 | 0.098 | 0.100 | 0.101 |
| *Enterocloster clostridioformis* YL32 | 0.500 | 0.523 | 0.516 | 0.513 | 0.136 | 0.119 | 0.148 | 0.134 |
| *Limosilactobacillus reuteri* | 0.049 | 0.058 | 0.058 | 0.055 | 0.042 | 0.026 | 0.031 | 0.033 |
| *Bifidobacterium animalis* | 0.111 | 0.111 | 0.111 | 0.111 | 0.027 | 0.014 | 0.002 | 0.014 |
| Total Euclidean distance | 0.572 | 0.599 | 0.59 | 0.587 | 0.385 | 0.336 | 0.351 | 0.357 |

### Benchmarking against Illumina short-read 16S rRNA amplicon sequencing

Illumina short-read sequencing has become established as a gold standard for 16S rRNA profiling^57,58^. As a result, there have been significant advancements in the algorithms and tools for classifying, analyzing, and inferring compositional and functional complexity from short-read amplicon sequencing data^52,59,60^. However, the short read length of Illumina reads restricts classification of 16S rRNA amplicons to the taxonomic levels of genus and family. Conversely, as our understanding of bacterial strain-specific metabolism and characteristics advances, there is a growing need for rapid and accurate classification from amplicon sequencing, ideally down to the strain level, to draw meaningful inferences for microbiota composition.

To benchmark our method against state-of-the-art Illumina 16S rRNA amplicon short-read sequencing, we compared the classification obtained with Illumina short-read sequencing data to the long-read nanopore sequencing data. We extracted DNA from intestinal contents of five unmanipulated OligoMM12 mice and sequenced the 16S rRNA V3-V4 hypervariable regions using Illumina short-read sequencing. From the same DNA samples, we amplified full-length 16S rRNA gene sequences using primers 27Fdeg/1492Rdeg and conducted long-read nanopore sequencing as described in the previous section. Classification against the non-redundant RDP database^52^ revealed that both methods yielded similar results regarding overall microbiota composition but with several key differences (**Figure 3**).

**Figure 3.**
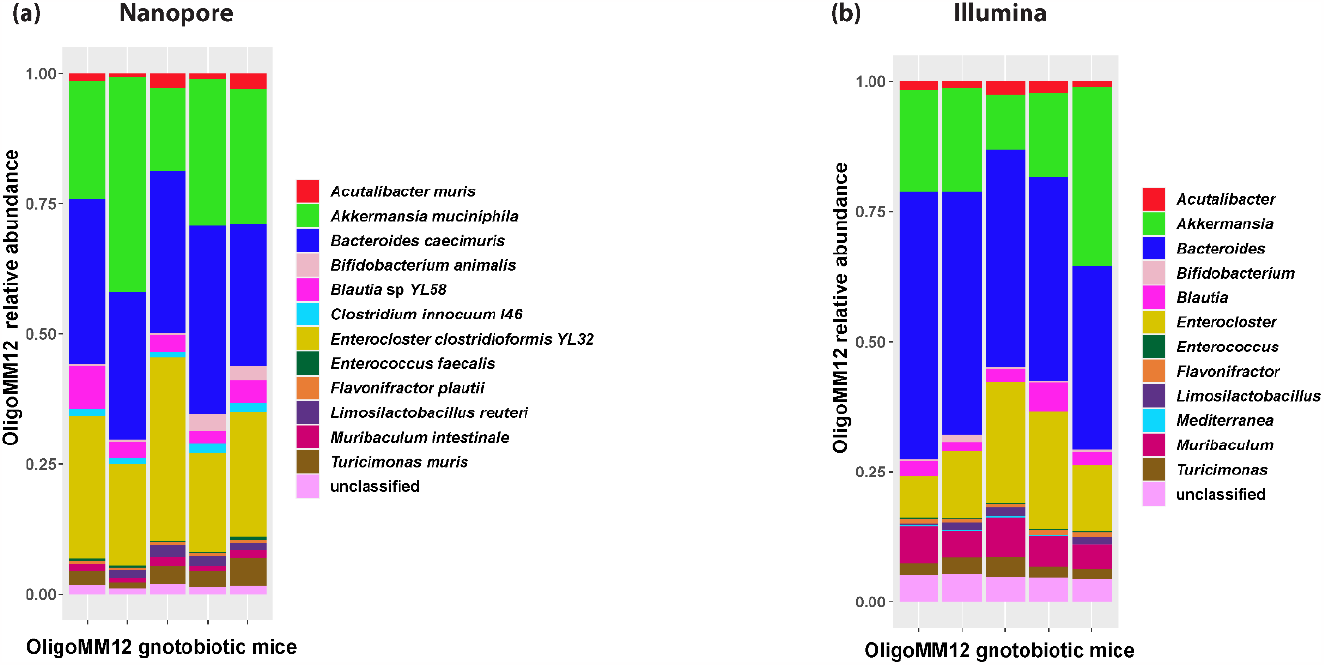
Benchmarking of full-length ONT nanopore against V3-V4 Illumina 16S rRNA gene amplicon sequencing. **(a)** Amplification by revised 27Fdeg-1492Rdeg primers and barcoding by SQK-NBD104/SQK-NB114 + SQK-LSK109 kit) performs better than **(b)** Illumina V3-V4 amplicon sequencing approach. Legends indicate the taxonomic levels (genus, species and strain) to which the reads could be classified. Reads with strain level classification are indicated with strain numbers (KB18, YL58, YL45, I46, YL32) in the legend. Each stacked bar represents one mouse individual and corresponds in both panels.

Notably, whilst classification of Illumina short-read data remained limited to the genus level, ONT full-length amplicon sequencing analysis achieved species-to strain-level resolution. We also observed that Illumina sequence data analysis yielded higher relative abundances of *Bacteroides* and *Muribaculum* compared to ONT sequencing. Furthermore, some of the Illumina reads were misclassified as belonging to the Bacteroidaceae genus *Mediterranea*. We also noticed that no taxon corresponding to *Clostridium innocuum* strain I46 was detectable in the Illumina sequencing results.

## Discussion

In this study, we systematically analyzed different library preparation protocols offered by ONT and demonstrated that a native barcoding approach (SQK-NBD104/SQK-NBD114), combined with customized primer pairs, outperforms the ONT proprietary 16S sequencing kit (SQK-16S024) that utilizes universal primers 27F/1492R. While the former is more time-intensive, the latter offers a streamlined single-step PCR approach to amplify and barcode the 16S rRNA amplicon. Classification accuracy is a crucial factor in microbial composition analysis, especially in the context of defined experimental low-complexity microbiota such as in gnotobiotic mice. For this application, our cautiously optimized native barcoding approach consistently provides more accurate results when compared to SQK-16S024.

We illustrated that appropriate primer design is paramount for 16S rRNA amplicon sequencing, more so for low-complexity bacterial communities and for detecting low-abundance bacteria. We found that our approach (with improved primer pair 27Fdeg/1492Rdeg) is robust in detecting low-abundance bacteria like *Bifidobacterium*.

Arguably, with increased microbiota species diversity it would become increasingly difficult to decrease analytical bias by optimizing 16S rRNA primer design. Thus, whilst for different low-complexity communities of known strain composition and 16S rRNA gene sequences, is doable and recommended to reduce analytical bias by rational primer optimization, we advise cautious use of the described primers 27Fdeg/1492Rdeg for the study of different low-complexity communities or high-complexity microbiota.

Species-specific quantitative 16S rRNA PCR has been developed as an alternative approach for similar applications. This more targeted and quantitative method can further decrease bias and sensitivity but is more difficult to establish and validate. Additionally, unlike untargeted 16S amplicon sequencing, it cannot be used to control for microbial contamination with unknown species.

We observed that in our experience the turnaround time taken of ONT sequencing, from the extraction of DNA to completion of microbiota compositional profiling, took about 3 days compared to typically ≥ 15 days for Illumina sequencing. While the preference for Illumina short-read versus ONT long-read sequencing depends on the scientific question and data analysis expertise, ONT sequencing offers multiple advantages including accessibility, shorter run time and affordability.

The pipeline described in this paper gave more intuitive results outperforming standard Illumina short-read or ONT 16S rRNA sequencing protocols, yet it has also some limitations. While the mock community of 9 bacteria has demonstrated our ONT sequencing approach is sufficiently accurate for characterizing a typical gnotobiotic gut community, benchmarking against other long-read sequencing methods will be required to evaluate the impact of sequencing errors introduced by ONT technology.

## Acknowledgements

We thank Alban Ramette, Miguel Terrazos, Stefan Neuenschwander and Sonja Gempler (Bioinformatics and Biostatistics group, IFIK, University of Bern) for helping with nanopore sequencing and data transfer, and Matheus Notter Dias (IFIK, University of Bern) for reisolating Oligo-MM12 bacterial strains. We thank staff and management team of the Clean Mouse Facility (CMF), University of Bern, for gnotobiotic animal maintenance and services. Data processing was performed on the IFIK-NGS Research Server and UBELIX (http://www.id.unibe.ch/hpc) HPC cluster at the University of Bern. This work was funded in part by the Swiss National Science Foundation (www.snf.ch; NCCR Microbiomes (grant 180575) and Sinergia grant 180317) and a grant from the Helmut-Horten-Stiftung (helmuthorten-stiftung.org; project “Synthetic Pathogen-Microbiota Symbiosis”)

